# PhageAI - Bacteriophage Life Cycle Recognition with Machine Learning and Natural Language Processing

**DOI:** 10.1101/2020.07.11.198606

**Authors:** Piotr Tynecki, Arkadiusz Guziński, Joanna Kazimierczak, Michał Jadczuk, Jarosław Dastych, Agnieszka Onisko

**Affiliations:** Computer Science Faculty of Bialystok University of Technology, Wiejska 45 A Street, 15-351 Bialystok; Proteon Pharmaceuticals SA, 90-364 Lodz, Poland

**Keywords:** PhageAI, bacteriophages, lifecycle, virulent, temperate, Machine Learning, Natural Language Processing, classifier, DNA embedding

## Abstract

**Background:** As antibiotic resistance is becoming a major problem nowadays in a treatment of infections, bacteriophages (also known as phages) seem to be an alternative. However, to be used in a therapy, their life cycle should be strictly lytic. With the growing popularity of Next Generation Sequencing (NGS) technology, it is possible to gain such information from the genome sequence. A number of tools are available which help to define phage life cycle. However, there is still no unanimous way to deal with this problem, especially in the absence of well-defined open reading frames. To overcome this limitation, a new tool is definitely needed.

**Results:** We developed a novel tool, called PhageAI, that allows to access more than 10 000 publicly available bacteriophages and differentiate between their major types of life cycles: lytic and lysogenic. The tool included life cycle classifier which achieved 98.90% accuracy on a validation set and 97.18% average accuracy on a test set. We adopted nucleotide sequences embedding based on the Word2Vec with Ship-gram model and linear Support Vector Machine with 10-fold cross-validation for supervised classification. PhageAI is free of charge and it is available at https://phage.ai/. PhageAI is a REST web service and available as Python package.

**Conclusions:** Machine learning and Natural Language Processing allows to extract information from bacteriophages nucleotide sequences for lifecycle prediction tasks. The PhageAI tool classifies phages into either virulent or temperate with a higher accuracy than any existing methods and shares interactive 3D visualization to help interpreting model classification results.

## Background

We are might soon be living in a post-antibiotic era and there is a need to find an alternative to treat microbial diseases, especially because of growing resistance of pathogens. One of the solutions that brings much scientific attention during recent years is phage therapy [1]. Bacteriophages are defined as viruses that target, infect, and replicate within bacteria, having high specificity restricted to one bacterial genus or even certain strains. They are among the most abundant entities on Earth - it is estimated that there are 10^31^ phages worldwide [2,3].

After their discovery at the beginning of the 20th century, phages suddenly lost popularity because antibiotics were discovered in parallel. Therefore, there are still many gaps in a knowledge of their biology. One of the problems that still needs to be addressed and investigated is a differentiation between the phage life cycles: lytic or lysogenic [3].) A virulent phage exhibits a strictly lytic life cycle in which after a phage attachment to a host cell, a nucleic acid is injected in order to use bacterial metabolism, replicate its genome and synthesize new virions. As a result, bacterium is lysed and bacteriophages are released to the environment. In contrast, a temperate phage carries a lysogenic cycle in which its genome might be inserted into a bacterial chromosome and form a prophage, a state in which it can last for many generations. However, when such a phage is induced with a certain stress factor, it can also enter a lytic cycle [3,4].

There is a need to define a life cycle of a phage especially when choosing phages for therapeutic purposes, as temperate phages are known to take part in a horizontal gene transfer (HTG). Since they can integrate into bacterial genomes, they can transfer undesirable features such as virulence factors or antibiotic resistance genes into subsequent bacterial generations. On the other hand, virulent phages are considered safe and are approved for use in phage therapy [2,5].

So far, there is no unambiguous and indisputable way to define a bacteriophage life cycle. There is a traditional experimental method based on clearance or turbidity of plaques, however, it is not of much use nowadays [5]. As NGS sequencing is less and less expensive, it becomes available for research units to gain information from bacteriophages’ sequences. In Andrew Millard lab webpage there is a plot presenting a cumulative number of phage genomes over the years (Figure 1). However, the analysis of phage sequences is still a struggle for the scientific community because of a low availability of reference genomes, mistakes in Open Reading Frames (ORFs) sizes done with automatic annotation programs and little knowledge about protein function of an analysed phage. Therefore, there are various approaches to define a phage life cycle [6,7]. It often starts with a search of reference sequences in Basic Local Alignment Search Tool (BLAST) and both automatic and manual annotation of genomes (e.g.in DNAMaster, University of Pittsburgh). Then, careful analysis of ORFs is done looking not only for sequence homology, but also for a structural one e.g.in HHpred and search for domains is performed in InterPro [8,9]. As a complement, phylogenetic analysis e.g., in MEGAX [10] and analysis of termini of phages in PhageTerm [11] are prepared.

**Figure 1.**
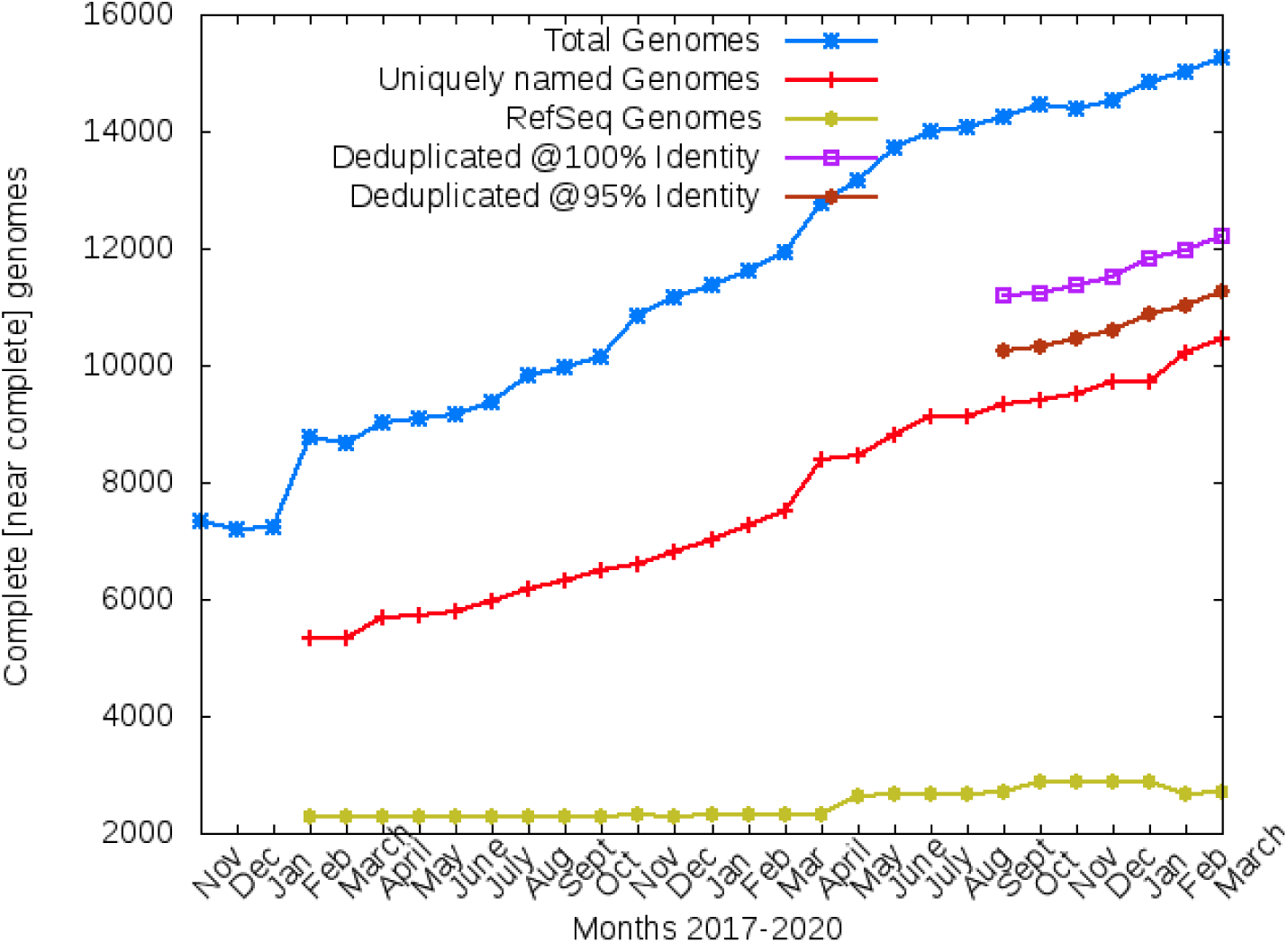
Number of sequenced phage genomes over the years. Published within author permission [13].

Currently, there is only one automatic tool called PHACTS in which a prediction of a phage life cycle is generated based on amino acid sequences of the analyzed phages. However, it requires an amino acid sequence based on annotation which can be imperfect and, moreover, it gives quite often averaged probability of results around 0.5 – 0.6 which is not satisfactory [12].

Consequently, there is a need to develop a fast and reliable tool which will be based on a phage genome analysis itself and which will not be dependent on hypothetical functions of potential ORFs which is very often the case for bacteriophage genomes with no reference. This is why we decided to apply solutions from the Artificial Intelligence (AI) domain that focuses on statistical models and algorithms allowing computer systems to solve a particular problem or perform a specific task with or without explicit expert rules and programmed instructions. While historically somewhat niche, increasingly better hardware and recent developments in the Deep Learning algorithms enable Machine Learning (ML) models and natural Language Processing (NLP) to achieve human-like performance on various tasks from multiple fields such as computer vision or knowledge extraction from the data. ML is applied to an increasingly wide range of domains. Every scientist has an opportunity to integrate it into his operations to become more competitive by gaining predictive insights and the potential to automate numerous tasks. Today’s AI frameworks are already mature and effective enough to be powerful tools not only for researchers but also for practical application developers.

In this paper, we present a novel approach based on Machine Learning and Natural Language Processing to classify phages into virulent or temperate based on their nucleotide sequences. Our tool is available online at https://phage.ai/.

## Results

This work was focused on constructing a novel Machine Learning and Natural Language Processing pipeline for bacteriophages’ life cycle classification. For this research we used 278 virulent and 174 temperate phage genomes in FASTA format.

We applied common NLP techniques for efficient DNA word embedding by k-mer structure (contiguous subsequence of *k* letters) with sliding window approach using constant *k = 6* and the Word2Vec algorithm with the Skip-gram model. It allowed us to extract vocabulary size *V = 4,096* used to represent each phage. The word embedding model was trained using sparse DNA 6-mers and produced dense embedding vectors consisting of 300 fixed-size numeric vector space. To maximize the performance of the algorithm we trained 20 iterations (epochs) and optimized neural network with negative sampling instead of hierarchy softmax function. The Word2Vec training setup parameters that we have used: *{size = 300, window = 5, min_count = 1, sample = 1e-3, sg = 1, hs = 0, iterations = 20, negative = 5, word_ngrams = 1, random_state = 42}*.

The final step for feature engineering was their selection. 150 nominal features were empirically chosen from a total of 300 vector size (Figure 2) by feature selection algorithm called Feature ranking with recursive feature elimination and cross-validated selection of the best number of features (RFECV) [14] using Support Vector Machine (SVM) estimator.

**Figure 2.**
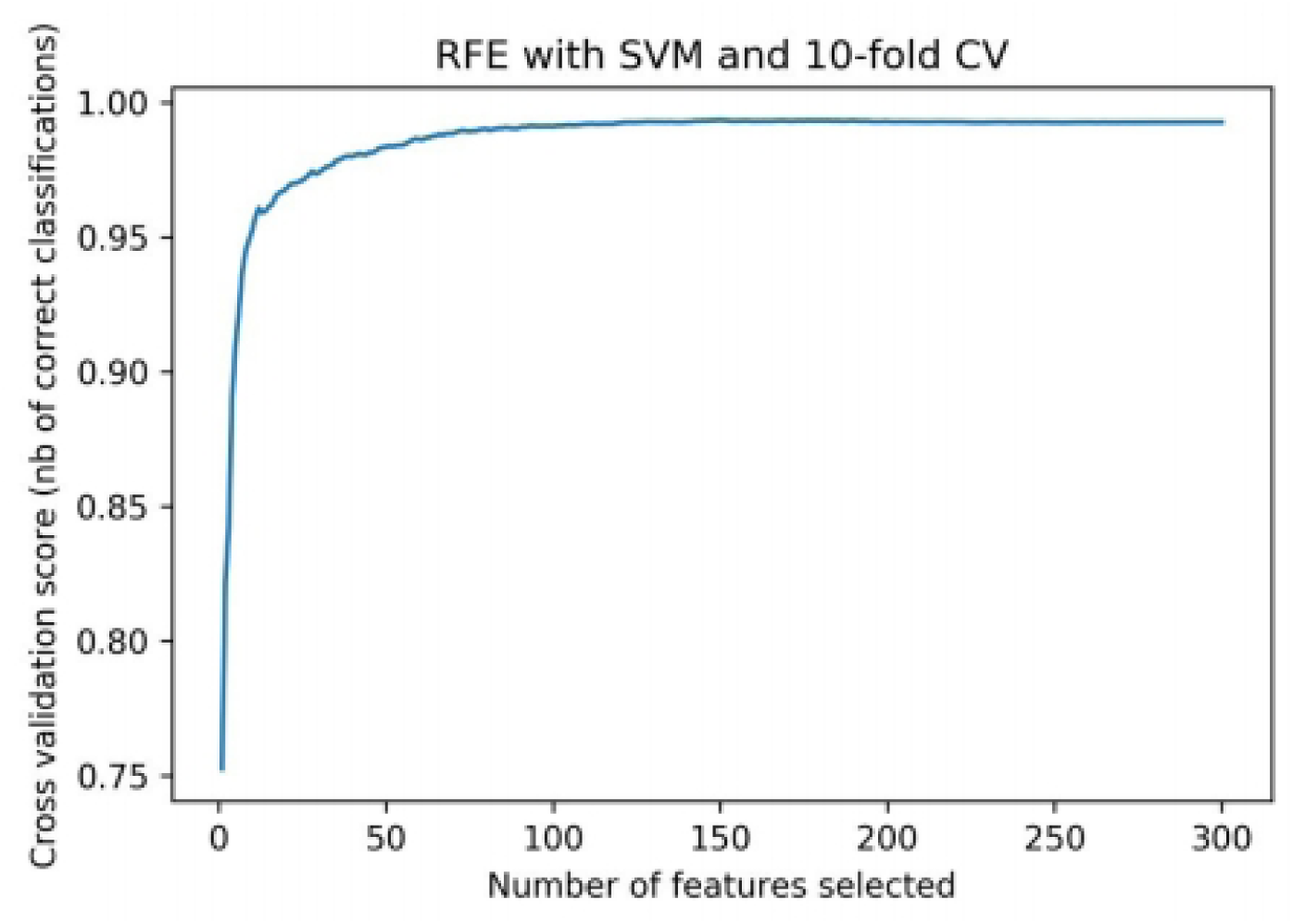
Optimal number of features for ROC AUC scoring

This number of features allowed us to train and tune 11 supervised ML classifiers (see Table 1). For tuning models hyperparameters, Bayesian optimization was applied with 10 fold cross-validation and F1-weighted scoring. The best result was achieved with a Support Vector Machine classifier with a linear kernel which resulted in an average accuracy of 98.90% on the validation sets (Table 2). The SVM training setup parameters that we have used after final tuning procedure: *{C = 1340*.*98, cache_size = 4000, class_weight = ‘balanced’, degree = 100, gamma = 1000, kernel = ‘linear’, probability = True, shrinking = False, random_state = 42}*.

**Table 1.** Bacteriophages life cycle prediction benchmark for 11 supervised ML tuned classifiers with 10-fold cross-validation

|  |  |  |  |  |  |
| --- | --- | --- | --- | --- | --- |
| MultinomialNB | 83.53 | 80.79 | 0.85 | 0.84 | 0.82 |
| SGDClassifier | 87.06 | 89.01 | 0.88 | 0.87 | 0.87 |
| MLPClassifier | 92.35 | 94.40 | 0.93 | 0.92 | 0.92 |
| LogisticRegression | 92.94 | 94.53 | 0.93 | 0.93 | 0.93 |
| RandomForestClassifier | 92.35 | 97.42 | 0.92 | 0.92 | 0.92 |
| **Support Vector Classification** | **98.90** | **99.84** | **0.99** | **0.99** | **0.99** |
| KNeighborsClassifier | 95.44 | 96.18 | 0.94 | 0.95 | 0.95 |
| GradientBoostingClassifier | 87.06 | 88.36 | 0.87 | 0.87 | 0.87 |
| XGBoost | 97.80 | 98.90 | 0.98 | 0.98 | 0.98 |
| CatBoostClassifier | 91.18 | 96.03 | 0.91 | 0.91 | 0.91 |
| LightGBM | 96.90 | 97.68 | 0.97 | 0.97 | 0.97 |
| **Classifier implementation name** | **Accuracy (%)** | **AUC (%)** | **Precision** | **Recall** | **F1-score** |

**Table 2.** The SVM model classification results on validation set

| Table 2. The SVM model classification results on validation set |  |  |  |  |
| --- | --- | --- | --- | --- |
| **Training set score** | 99.17% |  |  |  |
| **Validation set score** | 98.90% |  |  |  |
| **AUC** | 99.63% |  |  |  |
| **virulent** | 1.00 | 0.98 | 0.99 | 112 |
| **temperate** | 0.97 | 1.00 | 0.99 | 70 |
| **accuracy** |  |  | 0.99 | 182 |
| **macro avg** | 0.99 | 0.99 | 0.99 | 182 |
| **weighted avg** | 0.99 | 0.99 | 0.99 | 182 |
|  | **precision** | **recall** | **f1-score** | **phages** |

In-depth evaluation of the Support Vector Machine includes accuracy, precision, recall, F1-score and Area Under the Receiver Operating Characteristic (AUC) metrics as well as three plots: learning curve (Figure 3) which determines cross-validated training and test scores for different training set sizes, confusion matrix (Figure 4) for classification evaluation accuracy by computing the confusion matrix with each row corresponding to the true class and ROC AUC (Figure 5) which illustrates the diagnostic ability of a binary classifier system as its discrimination threshold is varied.

**Figure 3.**
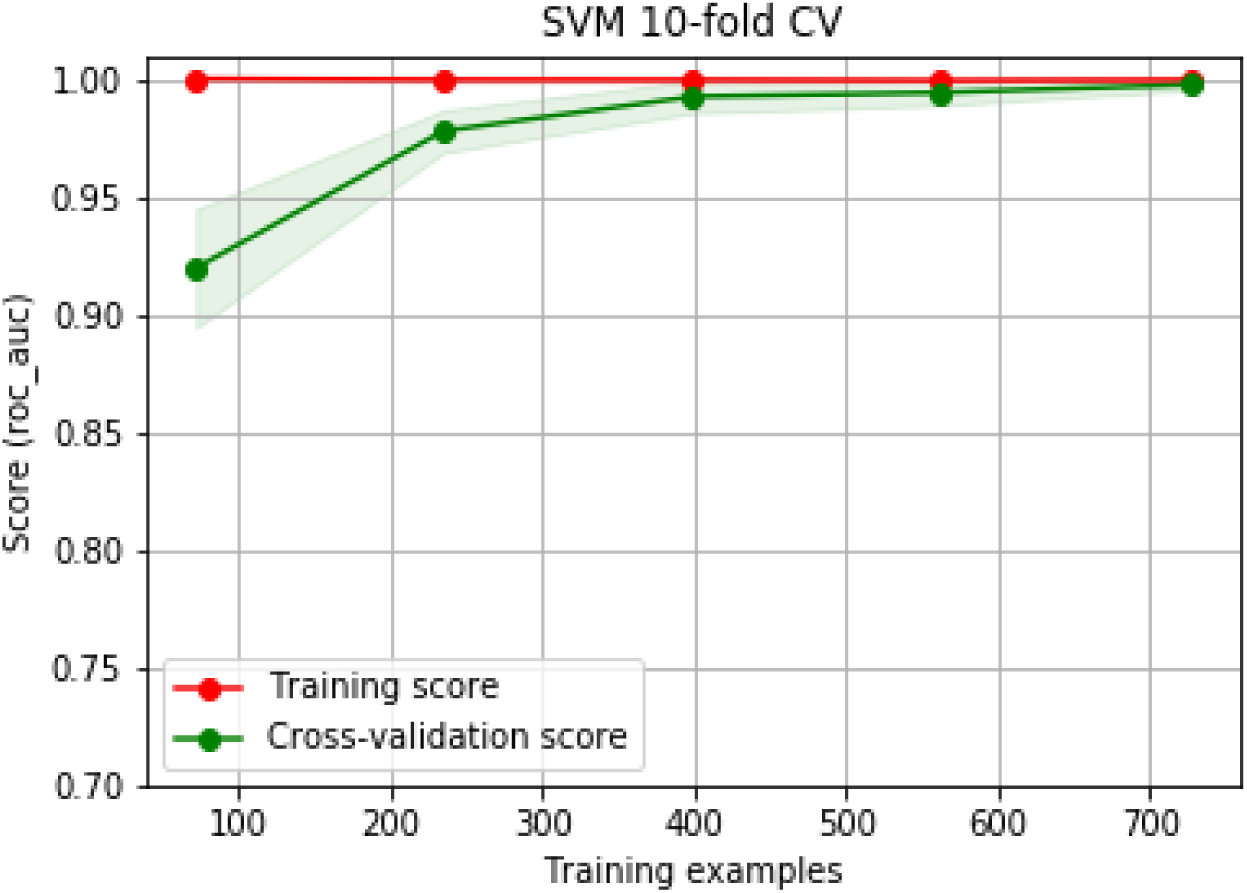
Learning curve with 10-fold CV for SVM on training dataset

**Figure 4.**
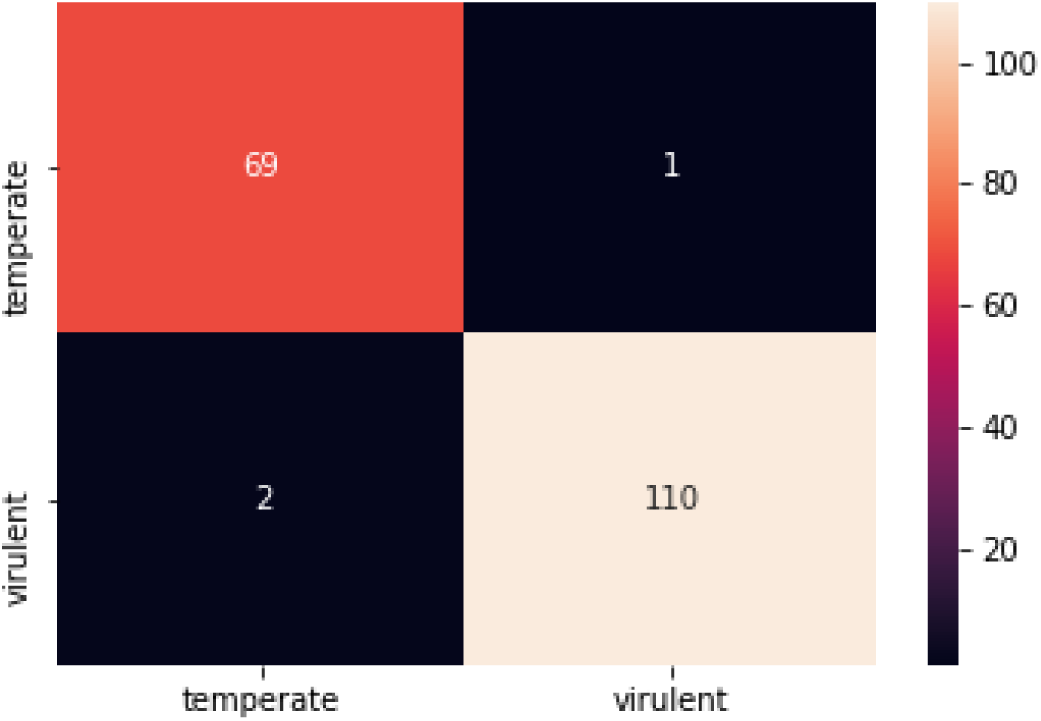
Confusion matrix for SVM on validation dataset

**Figure 5.**
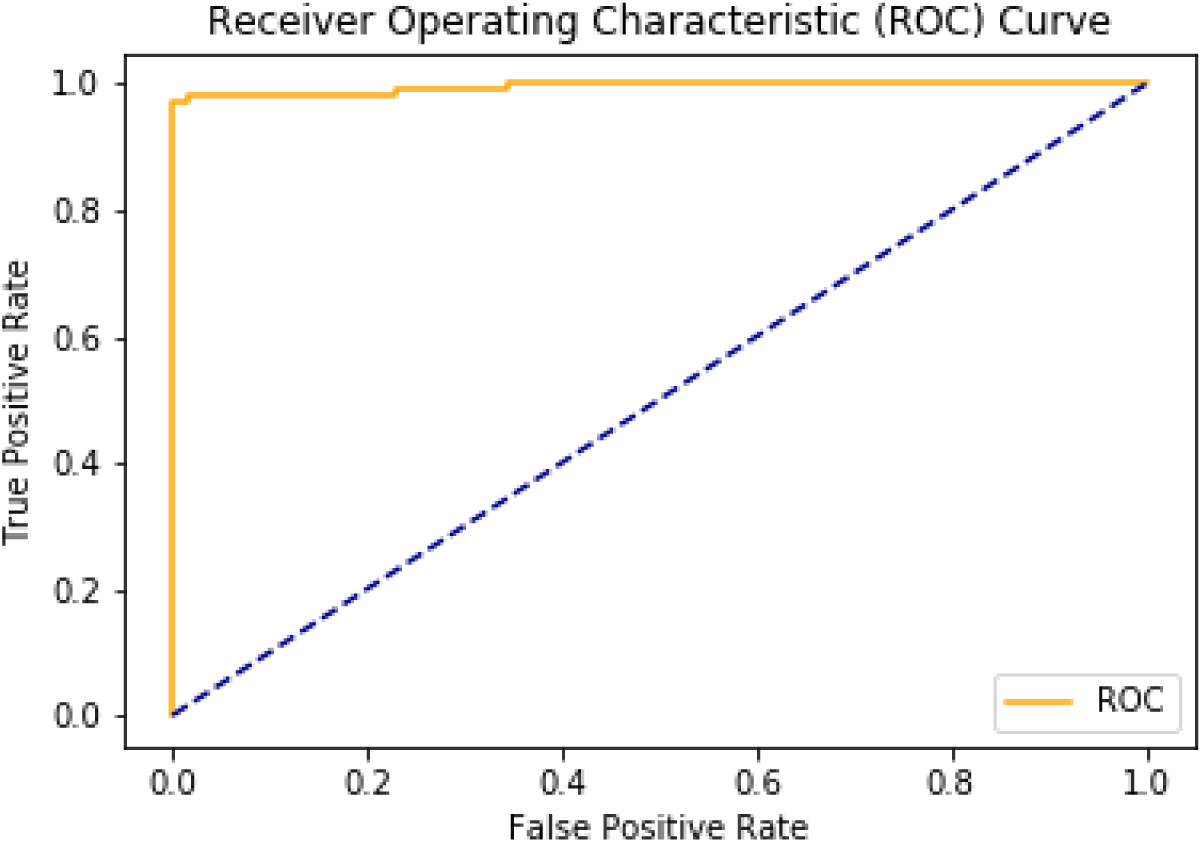
ROC AUC for SVM on validation dataset

The life cycles of all unseen bacteriophages from the testing dataset (54 virulent and 30 temperate phages from different species and families) were predicted correctly by the best SVM model, with 97.18% average accuracy across all classifiers. To confirm the model’s ability to generalize new data, we also tested it on a not publicly available dataset provided by the Proteon Pharmaceuticals S.A. company. All of 61 bacteriophages (49 virulent, 12 temperate) achieved correct prediction by the model, according to manual lifecycle prediction.

## Discussion

This study demonstrated that the PhageAI tool can automate bacteriophages life cycle classification. Nucleotide sequences in FASTA format are the only requirement to classify phages into virulent and temperate with accurate results. The PhageAI tool is free of charge, is available at https://phage.ai/ and can be adopted as a part of other custom bioinformatic pipelines using the available REST API. The PhageAI pipeline is flexible and susceptible to modification by new strategies, setups and frameworks on each of five existing steps (Figure 6) which opens up possibilities for custom adjustments.

**Figure 6.**
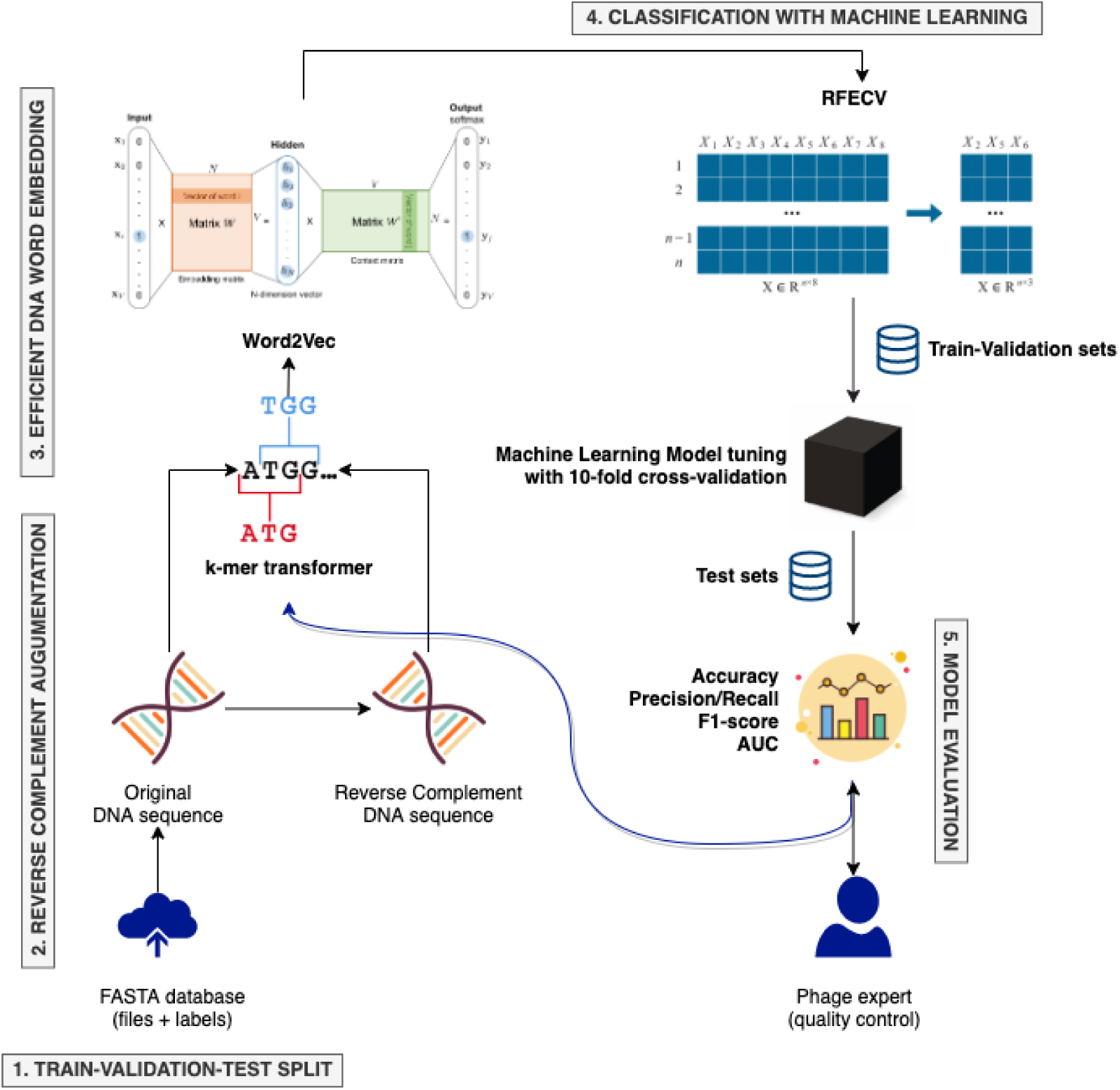
The *PhageAI* pipeline: proposed methodology for Bacteriophages life cycle recognition research

### Limitations and future directions

Interpretation of the SVM model is difficult given the multidimensional context of the data as well as the embedding used to get numeric vector space. Therefore, in the PhageAI tool we have prepared an interactive 3D visualization to help interpreting model classification results (Figure 7).

**Figure 7.**
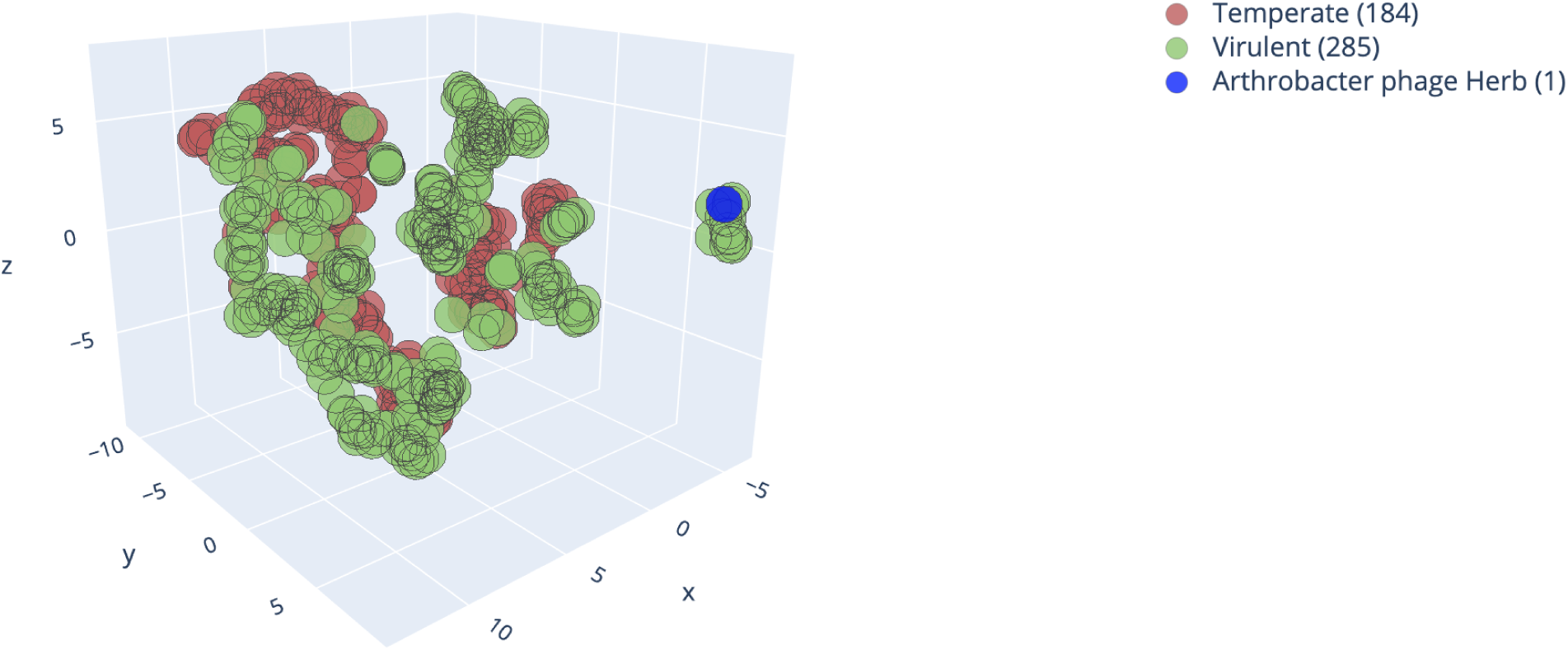
Virulent and temperate bacteriophages visualization via dimensionality reduction by UMAP.

The application of Uniform Manifold Approximation and Projection for Dimension Reduction (UMAP) [15] allowed us to separate between virulent and temperate phages and group a significant part of them into clusters and subgroups represented by the same life cycle. Therefore, as the next step of our research we intend to investigate this and rather look for a correlation between them.

The PhageAI tool is in active development. We also shared dedicated Python programming package *phageai* [16] and scheduled next life cycle classifier re-training sessions including new samples from different families and species and extended chronic infection prediction for filamentous bacteriophages.

### Review of existing solutions

Before developing the PhageAI tool, we have reviewed the existing solutions for bacteriophage life cycle classification. Namely, we focused on the tool that is currently the most popular and widely used for phage research: PHACTS [12]. It uses an ensemble of Random Forest classifiers trained on samples from the PHANTOME database [17]. The models use protein-based features representing calculated similarities to the analyzed genomes. However, since the proteins used by the classifiers are chosen mostly at random, the results vary greatly during practical tests, with the same phages classified as both virulent and temperate on multiple runs. By analyzing the entire nucleotide sequences and using techniques such as the reverse complement described in the methodology section, our approach yields much more stable results.

We also tested Virus Identification By iteRative ANnoTation (VIBRANT v1.2.1) software [18]. The tool utilizes a hybrid Machine Learning and protein similarity approach that is not reliant on sequence features for automated recovery and annotation of viruses, determination of genome quality and completeness, and characterization of virome function from metagenomic assemblies. VIBRANT uses supervised neural network Multi-layer Perceptron classifier (MLP) with protein signatures and a custom v-score metric that circumvents traditional boundaries to maximize identification of lytic viral genomes and integrated proviruses, including highly diverse viruses. Surprisingly, during testing of Enterobacteria phage Mu (NC_000929.1), which is a model example of phage integrating into a host genome using the transposition process [19], the program indicated that it has a virulent character. Therefore, one has to be cautious when relaying on the life cycle assessment presented by the program. At the same time, this program is an ideal tool for rapid annotation of the viral genome, which allows manual review of the program’s indications.

## Conclusions

In this paper, we have shown that it is possible to capture and extract knowledge from hundreds of bacteriophages sequences to classify their life cycles with a high accuracy and immediate result. The PhageAI tool needs only DNA nucleotide sequence in FASTA format to make a prediction, which is to our best knowledge a novel approach.

The application of Machine Learning and Natural Language Processing for bacteriophage research allowed to distinguish between the two major types of their life cycles: lytic and lysogenic. Furthermore, it achieved accurate results on unknown phages which was double tested on unseen data.

Currently adopted methodology opens up opportunities for further research in the field of phage classification. Extending the PhageAI tool with DNA word embedding like transfer-learning or context-based bidirectional models might increase sequence-based classification performance for other issues such as predicting proteins features, distinguishing bacteriophages taxonomy or phage host range identification. Deep Learning approach is becoming more justified in the next step because the PhageAI repository has already collected more than 10,000 phages’ sequences.

PhageAI was released as a free web platform, REST API service and open sourced Python package which should allow other researchers to include our tool in their pipelines.

## Methods

In our study we have used more than 600 genomic sequences of bacteriophages from ACLAME [20] and PhagesDB [21] with information about their life cycle. We manually verified predictions for the purpose of this study. In order to standardize the annotation, all phage genomes were annotated by using DNAMaster (a tool developed by Dr. Jeffrey Lawrence, the University of Pittsburgh, v5.23.3) with its auto annotations option which combines Glimmer [22] and GenMarkS [23] algorithms. Then, all detected ORF were analyzed to find proteins that may be involved in bacteriophage integration into the host genome. For this purpose, HMMscan [24] and InterPro ([9] access: 07.2019) software were used, which allow for detection of characteristic domains in the protein sequence. Additionally, Hhpred ([8] access: 07.2019) was run to find remote homologs based on the modeled 3D structure. In the case when none of lysogenic factors was found or a phage was unable to maintain the lysogeny, the phage sequence was marked as virulent, otherwise it was temperate. The phages for which the life cycle could not be predicted or was unclear were discarded from further processing.

Moreover, amino acid sequences of phages were analyzed in the PHACTS tool to compare the predictions obtained manually.

### Datasets

Final training dataset after manual editing consisted of 278 virulent and 174 temperate phages. Additionally, we selected a testing dataset of 54 virulent and 30 temperate phages from different species and families (Figure 8, Figure 9, Figure 10).

**Figure 8.**
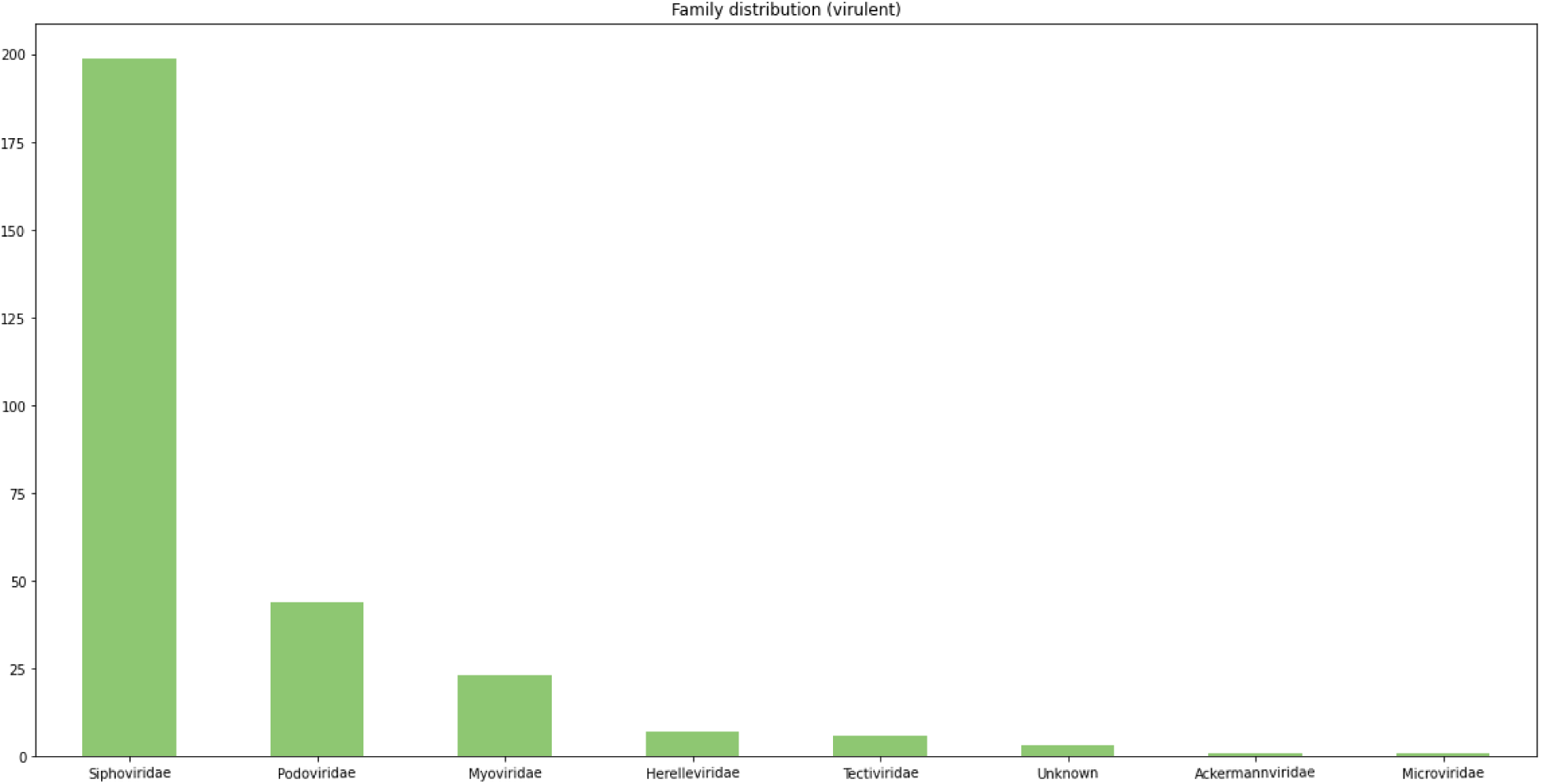
Training dataset family distribution for virulent phages

**Figure 9.**
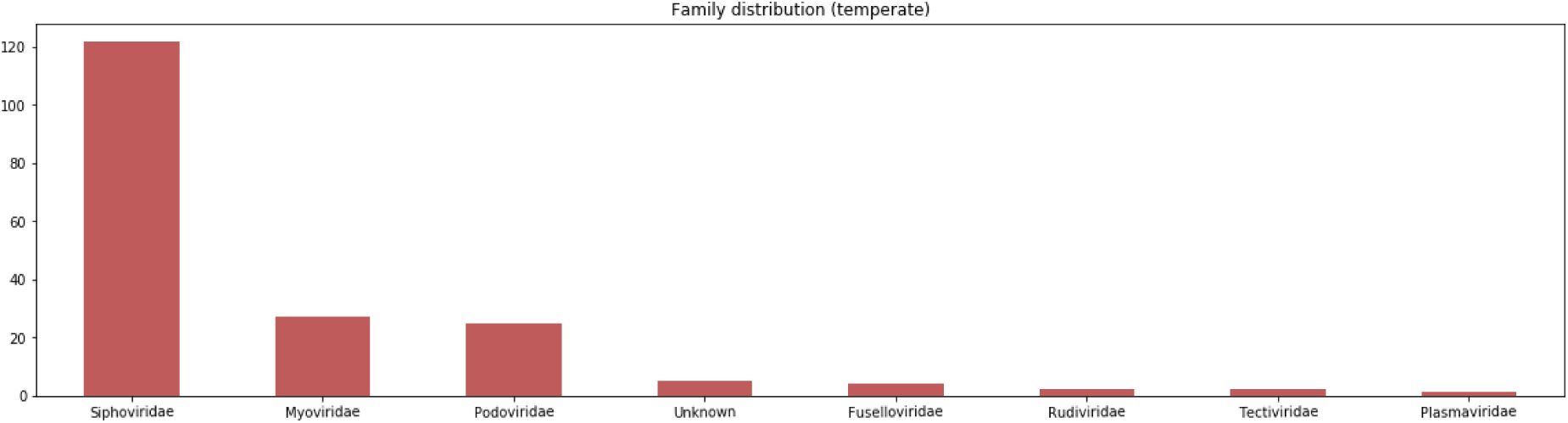
Training dataset family distribution for temperate phages

**Figure 10.**
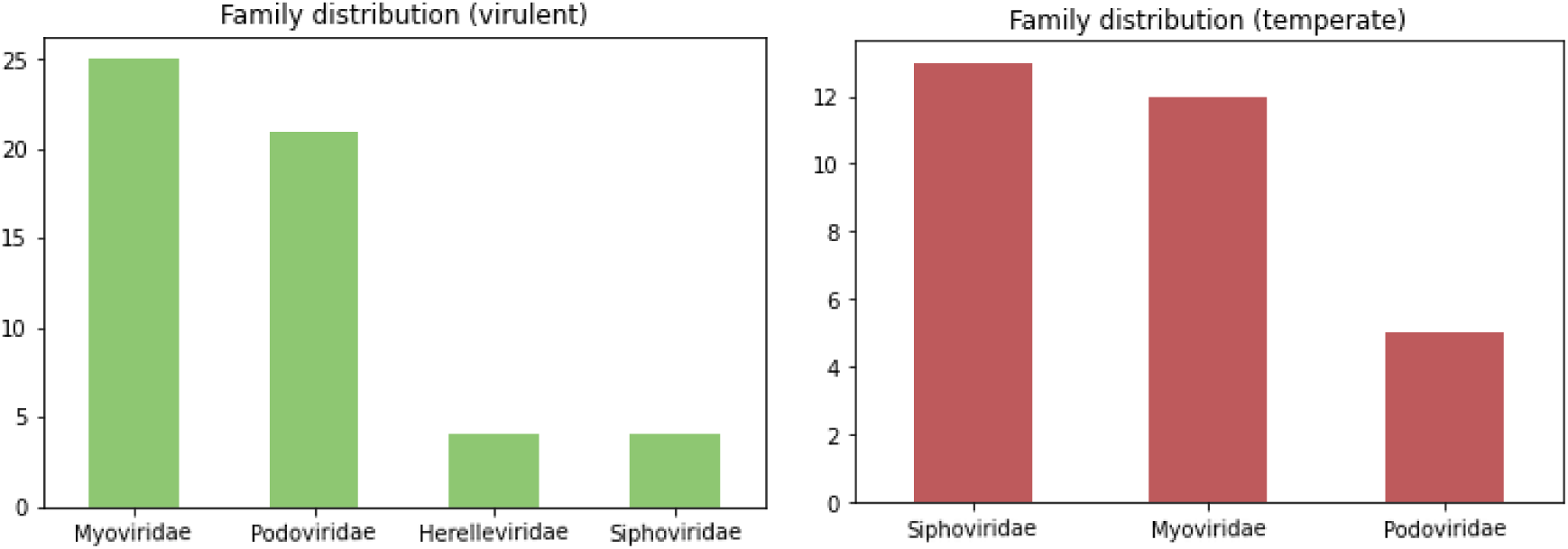
Testing dataset family distribution for virulent and temperate phages

All bacteriophages sequences with NCBI accession numbers are available in the Availability of Data and Materials section.

### The PhageAI pipeline

The PhageAI pipeline uses open source Python libraries: Biopython v1.76 [25], gensim v3.8.1 [26], scikit-learn v0.22.1 [27], xgboost v1.0.1 [28], catboost v0.22 [29], lightgbm v2.3.1 [30], scikit-optimize v0.7.4 [31] and matplotlib v3.2.0 [32].

The *PhageAI* pipeline covers 5 steps (Figure 6):

1. Train-validation-test split
2. Reverse complement augmentation
3. Efficient DNA word embedding
4. Classification with Machine Learning:
  a. Efficient feature selection
  b. Supervised learning
5. Model evaluation

### Train-validation-test split

To control how much a ML model is learning from the data, a well-established practice is to split samples to evaluate the future classifier with different bacteriophages. To train and evaluate the results we have chosen the following approaches:

- Cross-validation: stratified shuffle with 10-folds and 80% - 20% train-validation proportions was used to find the optimal hyperparameters and evaluate the results during training. For data stratification we used life cycles as well as bacteriophages families values to preserve the percentage of samples for each class.
- Holdout validation: a dataset of 84 unseen samples was designated as the testing set. It was not used in the training process directly, but it was employed to compare the model’s metrics after training.
- Additional holdout validation: the second dataset delivered by Proteon Pharmaceuticals S.A. company, containing 61 samples unavailable to the models during training was used to estimate the final metrics.

### Reverse complement augmentation

After train-validation-test split the reverse complement bacteriophage sequences (Figure 11) were treated as another samples. It enabled the ML model to automatically learn the complex relationships between the double strand DNA sequences.

**Figure 11.**
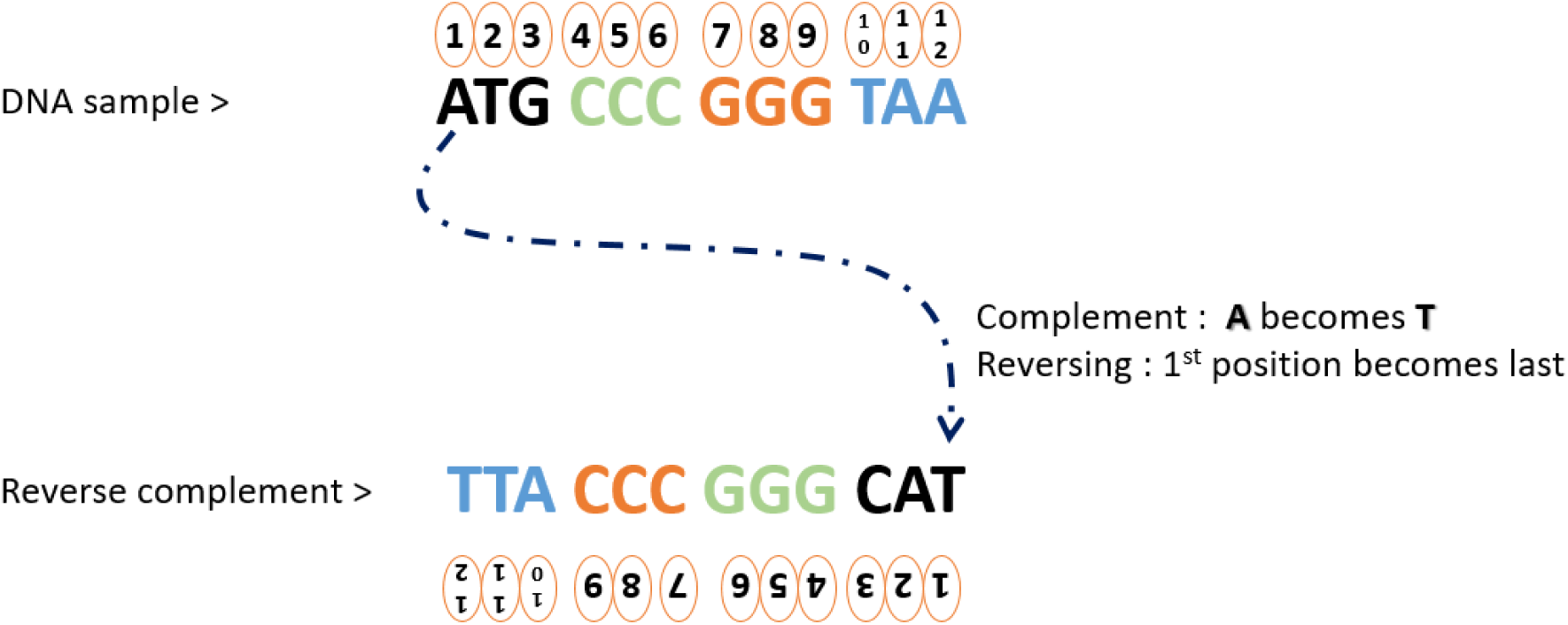
Reverse complement of a DNA sequence. Published within author permission [33].

Previous studies [34, 35] confirm the importance of utilizing the reverse complement DNA sequences, which is connected with data augmentation. This step also allowed us to double the datasets which became ready to be vectorized.

### Efficient DNA word embedding

Bacteriophages genomic sequences in FASTA format are represented as relatively long strings (between 5,000 - 300,000 bp) consisting of nucleotides *{A, C, G, T}*. This format of data is somewhat challenging for Machine Learning algorithms, where selection of a fixed-size feature list is the key to a robust classification model. A naive approach of transforming the whole DNA into a N-dimensional one-hot encoding space is very memory-consuming and devoid of the relevance of nucleotides in the genome. To build a representative vector space for bacteriophages sequences and drastically reduce memory requirements to accelerate further classification, we adopt common NLP techniques based on distributed representations of overlapping k-mers components and word embeddings.

The k-mer structure concept is commonly employed to represent long sequences of amino acids or nucleotides. For example, a 9-mer needs a vector of dimension 4^9^ = 262,144. The higher k-value could be problematic when applying the classification models to solve problems in DNA sequence analysis, especially since most of the ML algorithms prefer lower-dimensional continuous vectors as input. Therefore, we tested and compared three methods (sliding window, non-overlapping and variable-length) for k-mers extraction of length 3 <= k <= 12 and their impact on our experiment. Additionally, all the k-mers which contain characters outside of the nucleotide alphabet *{A, C, G, T}* were removed before vectorization was launched. This includes characters used to signify uncertainties within the sequenced genome.

One of the key ideas in NLP is how to efficiently convert sequences of character or words into numeric vectors, which then can be fed into various ML models. To obtain feature vectors of fixed size representing the genomes, we adopted word embedding based on the Word2Vec with Ship-gram model, which leverages a shallow neural network with a projection layer. The Skip-gram model is an efficient method trained to predict the probabilities of a word being a context word for the given target. The “context” is a set of adjacent subsequences surrounding the targeted k-mer. Using fixed length vectors to represent the sequence, the similarity between bacteriophages can be measured, even though each sequence can be of a different length (bp).

Finally, bacteriophages DNA were represented by the average of the k-mer embedding vectors of words that compose the sequences, which means that each genome was described by averaged numeric values in vector space. The idea of averaged word embeddings was adopted from X et al., 20xx where averaged word embeddings were used for document paragraphs [36].

### Classification with Machine Learning

#### Efficient feature selection

Heterogeneous features extracted from average of the k-mer embedding vectors might reflect better pattern information for characterizing bacteriophages lifecycle. For this purpose, we applied an RFECV which is an efficient feature selection method to remove irrelevant attributes and ceiling the generalization ability of the next step model.

#### Supervised learning

For this study we trained and compared results from 11 implementations of supervised ML algorithms:

- Bayesian models: *MultinomialNB*
- Support-vector machines: *SVC, SGDClassifier*
- Linear models: *LogisticRegression*
- Neural networks: *MLPClassifier*
- Decision trees: *RandomForestClassifier*
- Similarity-based algorithms: *KNeighborsClassifier*
- Gradient boosting: *GradientBoostingClassifier, XGBoost, CatBoostClassifier, LightGBM*.

For tuning models hyperparameters we discarded techniques such as Grid Search and Randomized Search which search through the entire space of available parameter combinations in an isolated way without improving based on the past results. Instead, we applied Bayesian Optimisation [37], which minimizes the time spent to obtain an optimized set of model parameters. We measured the accuracy, precision, recall, F1-score and Area Under the Receiver Operating Characteristic (AUC). To increase the performance of gradient-based classifiers we trained them with multiple NVIDIA GPUs usage.

## List of abbreviations

AUC: Area Under Curve
ML: Machine Learning
MLP: Multi-layer Perceptron classifier
NGS: Next Generation Sequencing
NLP: Natural Language Processing
RFECV: Feature ranking with recursive feature elimination and cross-validated selection of the best number of features
ROC AUC: area under an ROC curve
SVM: Support Vector Machine
UMAP: Uniform Manifold Approximation and Projection

## Declarations

### Ethics approval and consent to participate

Not applicable.

### Consent for publication

Not applicable.

### Availability of Data and Materials

The dataset used and analysed during the current study are available in the NCBI repository. https://pmlegacy.ncbi.nlm.nih.gov/sites/myncbi/1Xu-9lbsnfN1Wi/collections/59768502/public/

### Competing interests

In accordance with *PhageAI - Bacteriophage Life Cycle Recognition with Machine Learning and Natural Language Processing* policy, the authors are reporting that the PhageAI platform was developed by the authors and Proteon Pharmaceuticals is the owner of the platform.

### Funding

This work was supported by NVIDIA GPU Grant Program and by the Bialystok University of Technology research grant no. MB/WI/I/2018 funded by the Ministry of Science and Higher Education of Poland.

### Authors’ contributions

PT adopted and trained the Word2Vec and ML classifiers, implemented the PhageAI code base. AG and JK introduced domain knowledge about the phages, selected and manually verified samples included in train and test sets. MJ suggested the Word2Vec usage for DNA sequences embedding and evaluated the PHACTS methodology. PT, JK and AG prepared the manuscript. AO and JD conceptualized the study and revised the manuscript. All authors read and approved the final manuscript.

## Acknowledgements

The authors thank Sebastian Burzyński for his invaluable help with server and network infrastructure setup, Yana Minina for supporting NLP and ML classification by multiply setups preparation in the PhageAI pipeline and Mammal Research Institute Polish Academy of Sciences which shared GPU server for this research computation. Last, not least, thanks to all those who deposit their bacteriophages data in public databases, and to those who maintain these databases.

## Notes

https://phage.ai/

## References

[1] Lin DM, Koskella B, Lin HC. Phage therapy: An alternative to antibiotics in the age of multi-drug resistance. World J Gastrointest Pharmacol Ther. 2017;8(3):162.

[2] Jassim SAA, Limoges RG. Natural solution to antibiotic resistance: Bacteriophages “The Living Drugs.” Vol. 30, World Journal of Microbiology and Biotechnology. 2014. p. 2153–70.

[3] Doss J, Culbertson K, Hahn D, Camacho J, Barekzi N. A review of phage therapy against bacterial pathogens of aquatic and terrestrial organisms. Vol. 9, Viruses. 2017.

[4] Letchumanan V, Chan KG, Pusparajah P, Saokaew S, Duangjai A, Goh BH, et al. Insights into bacteriophage application in controlling vibrio species. Vol. 7, Frontiers in Microbiology. 2016.

[5] Hyman P. Phages for phage therapy: Isolation, characterization, and host range breadth. Vol. 12, Pharmaceuticals. 2019.

[6] Oduor JMO, Kiljunen S, Kadija E, Mureithi MW, Nyachieo A, Skurnik M. Genomic characterization of four novel Staphylococcus myoviruses. Arch Virol. 2019;164(8):2171–3.

[7] Kazimierczak J, Wójcik EA, Witaszewska J, Guziński A, Górecka E, Stańczyk M, et al. Complete genome sequences of Aeromonas and Pseudomonas phages as a supportive tool for development of antibacterial treatment in aquaculture. Virol J. 2019;16(1).

[8] Zimmermann L, Stephens A, Nam SZ, Rau D, Kübler J, Lozajic M, et al. A Completely Reimplemented MPI Bioinformatics Toolkit with a New HHpred Server at its Core. J Mol Biol. 2018;430(15):2237–43.

[9] Jones P, Binns D, Chang HY, Fraser M, Li W, McAnulla C, et al. InterProScan 5: Genome-scale protein function classification. Bioinformatics. 2014;30(9):1236–40.

[10] Kumar S, Stecher G, Li M, Knyaz C, Tamura K. MEGA X: Molecular evolutionary genetics analysis across computing platforms. Mol Biol Evol. 2018;35(6):1547–9.

[11] Garneau JR, Depardieu F, Fortier LC, Bikard D, Monot M. PhageTerm: A tool for fast and accurate determination of phage termini and packaging mechanism using next-generation sequencing data. Sci Rep. 2017;7(1).

[12] McNair K, Bailey BA, Edwards RA. PHACTS, a computational approach to classifying the lifestyle of phages. Bioinformatics. 2012;28(5):614–8.

[13] MillardLab website: http://millardlab.org/bioinformatics/bacteriophage-genomes/phage-genomes-march2020/, Accessed 24 April 2020.

[14] Mikolov T, Sutskever I, Chen K, Corrado G, Dean J. Distributed representations ofwords and phrases and their compositionality. In: Advances in Neural Information Processing Systems. 2013.

[15] McInnes L, Healy J, Saul N, Großberger L. UMAP: Uniform Manifold Approximation and Projection. J Open Source Softw. 2018.

[16] PhageAI tool as Python package: https://pypi.org/project/phageai/, Accessed 24 April 2020.

[17] The PhAnToMe database of over 1,000 phage genomes, http://www.phantome.org/, Accessed 24 April 2020.

[18] Kieft K, Zhou Z, Anantharaman K. VIBRANT: automated recovery, annotation and curation of microbial viruses, and evaluation of viral community function from genomic sequences. Microbiome. 2020.

[19] Harshey RM. Transposable Phage Mu. In: Mobile DNA III. 2015.

[20] Leplae R. ACLAME: A CLAssification of Mobile genetic Elements. Nucleic Acids Res. 2004.

[21] Russell DA, Hatfull GF. PhagesDB: The actinobacteriophage database. Bioinformatics. 2017.

[22] Delcher AL, Bratke KA, Powers EC, Salzberg SL. Identifying bacterial genes and endosymbiont DNA with Glimmer. Bioinformatics. 2007.

[23] Besemer J, Lomsadze A, Borodovsky M. GeneMarkS: A self-training method for prediction of gene starts in microbial genomes. Implications for finding sequence motifs in regulatory regions. Nucleic Acids Res. 2001.

[24] Potter SC, Luciani A, Eddy SR, Park Y, Lopez R, Finn RD. HMMER web server: 2018 update. Nucleic Acids Res. 2018.

[25] Biopython is a set of freely available tools for biological computation, https://biopython.org/, Accessed 24 April 2020.

[26] gensim is a software to realize unsupervised semantic modelling from plain text, https://radimrehurek.com/gensim/, Accessed 24 April 2020.

[27] scikit-learn Machine Learning package in Python, https://scikit-learn.org/, Accessed 24 April 2020.

[28] XGBoost is a scalable and flexible gradient boosting algorithm implementation in Python, https://xgboost.ai/, Accessed 24 April 2020.

[29] CatBoost is a high-performance open source library for gradient boosting on decision trees, https://catboost.ai/, Accessed 24 April 2020.

[30] LightGBM is a fast, distributed, high performance gradient boosting framework based on decision tree algorithms, used for ranking, classification and many other machine learning tasks, https://github.com/microsoft/LightGBM, Accessed 24 April 2020.

[31] Shcherbatyi I., Head T. and Louppe G., Scikit-learn hyperparameter search wrapper, https://scikit-optimize.github.io/, Accessed 24 April 2020.

[32] Matplotlib is a comprehensive library for creating static, animated, and interactive visualizations, https://matplotlib.org/, Accessed 24 April 2020.

[33] Shad Arf blog: https://shadarf.blogspot.com/2017/07/how-to-make-reverse-complement-of-dna.html, Accessed 24 April 2020.

[34] Cao Z, Zhang S. Simple tricks of convolutional neural network architectures improve DNA- protein binding prediction. Bioinformatics. 2019;

[35] Shrikumar A, Greenside P, Kundaje A, Science C. Reverse-complement parameter sharing improves deep learning models for genomics. BioRxiv. 2017;

[36] Andrew M. Dai. Document Embedding with Paragraph Vectors. Arxiv. 2015;

[37] Guyon I, Weston J, Barnhill S, Vapnik V. Gene selection for cancer classification using support vector machines. Mach Learn. 2002;

